# Analysis of KSHV B lymphocyte lineage tropism in human tonsil reveals efficient infection of CD138+ plasma cells

**DOI:** 10.1101/2020.01.17.910612

**Authors:** Farizeh Aalam, Romina Nabiee, Jesus Ramirez Castano, Jennifer E. Totonchy

## Abstract

Despite 25 years of research, the basic virology of Kaposi Sarcoma Herpesviruses (KSHV) in B lymphocytes remains poorly understood. This study seeks to fill critical gaps in our understanding by characterizing the B lymphocyte lineage-specific tropism of KSHV. Here, we use lymphocytes derived from 40 human tonsil specimens to determine the B lymphocyte lineages targeted by KSHV early during *de novo* infection in our *ex vivo* model system. We characterize the immunological diversity of our tonsil specimens and determine that overall susceptibility of tonsil lymphocytes to KSHV infection varies substantially between donors. We demonstrate that a variety of B lymphocyte subtypes are susceptible to KSHV infection and identify CD138+ plasma cells as a rare, but highly targeted cell type for *de novo* KSHV infection. Finally, we determine that the host immunological status influences the course of *de novo* infection in B lymphocytes. These results improve our understanding of KSHV transmission and the biology of early KSHV infection in a naïve human host, and lay a foundation for further characterization of KSHV molecular virology in B lymphocyte lineages.

## Introduction

Kaposi Sarcoma-associated Herpesvirus (KSHV/HHV-8) is a lymphotrophic gamma-herpesvirus. In addition to its role in the pathogenesis of Kaposi Sarcoma (KS) {Chang:1994up}, KSHV infection is associated with two lymphoproliferative disorders, multicentric Castleman disease (MCD) and primary effusion lymphoma (PEL) {Cesarman:1995ke, Soulier:1995tc}, as well as a recently characterized inflammatory disorder KSHV inflammatory cytokine syndrome (KICS) {Uldrick:2010gw}. Although KSHV-associated lymphoproliferative disorders are rare, their incidence has not declined as HIV treatment has improved {Powles:2009fo, Bhutani:2015fc} suggesting that, in contrast to KS, immune reconstitution is not sufficient to prevent KSHV-associated lymphoproliferative disease in people living with HIV/AIDS. Moreover, the KSHV-associated lymphoproliferative diseases are uniformly fatal with few effective treatment options {Carbone:2014ez}.

Despite the fact that KSHV is lymphotropic and causes pathological lymphoproliferation *in vivo*, study of *de novo* KSHV infection in B lymphocytes has historically been difficult {Kang:2017jq}. Resting peripheral B cells and many established B cell-derived cell lines are refractory to KSHV infection {Rappocciolo:2008gy}, but several groups have recently been successful in infecting B lymphocytes derived from human tonsil {Hassman:2011dj, Myoung:2011cn, Myoung:2011ir, Bekerman:2013hy, Totonchy:2018ir, Nicol:2016ga}. KSHV DNA is detectable in human saliva and salivary transmission is thought to be the primary route of person-to-person transmission for KSHV {Casper:2007dp, Pauk:2000cq, Casper:2004vz, Vieira:1997vi}, making the oral lymphoid tissues a likely site for the initial infection of B lymphocytes in a naïve human host. Thus, in addition to being susceptible to *ex vivo* infection, tonsil lymphocytes represent a highly relevant model for understanding early infection events in KSHV transmission.

The existing studies of KSHV infection in tonsil-derived B cells have explored a limited number of cell surface markers including IgM, immunoglobulin lambda light chains and activation markers on infected cells {Hassman:2011dj, Nicol:2016ga, Totonchy:2018ir}. However, no studies to date have comprehensively explored the specific B lymphocyte lineages targeted by KSHV infection in human tonsil specimens.

In this study, we performed KSHV infection of 40 human tonsil specimens from diverse donors and utilized lineage-defining immunological markers by flow cytometry to establish the primary B cell lineage tropism of KSHV. Our results demonstrate that the susceptibility of tonsil-derived B lymphocytes to *ex vivo* KHSV infection varies substantially from donor-to-donor, and that a variety of B cell lineages are susceptible to KSHV infection. In particular, CD138+ plasma cells are highly targeted by KSHV infection despite the fact that they are present at low frequencies in tonsil tissue. We demonstrate that high susceptibility of plasma cells to KSHV infection cannot entirely be explained by the presence of the CD138 heparin sulfate proteoglycan as an attachment factor. Finally, we demonstrate that although the baseline T cell microenvironment does not seem to influence susceptibility of tonsil specimens to KSHV infection, manipulation of CD4/CD8 T cell ratios can alter the targeting of specific B cell lineages by KSHV. These results provide new insights into early events driving the establishment of KSHV infection in the human immune system and demonstrate that alterations in immunological status can affect the dynamics of KSHV infection in B lymphocytes.

## Materials and Methods

### Reagents and Cell Lines

CDw32 L cells (CRL-10680) were obtained from ATCC and were cultured in DMEM supplemented with 20% FBS (Sigma Aldrich) and Penicililin/Streptomycin/L-glutamine (PSG/Corning). For preparation of feeder cells CDw32 L cells were trypsinized and resuspended in 15 ml of media in a petri dish and irradiated with 45 Gy of X-ray radiation using a Rad-Source (RS200) irradiator. Irradiated cells were then counted and cyropreserved until needed for experiments. Cell-free KSHV.219 virus derived from iSLK cells {Myoung:2011ic} was a gift from Javier G. Ogembo (City of Hope). Human tonsil specimens were obtained from NDRI (www.ndri.org). Human fibroblasts were derived from primary human tonsil tissue and immortalized using HPV E6/E7 lentivirus derived from PA317 LXSN 16E6E7 cells (ATCC CRL-2203). Antibodies for flow cytometry were from BD Biosciences and Biolegend and are detailed below.

### Isolation of primary lymphocytes from human tonsil

De-identified human tonsil specimens were obtained after routine tonsillectomy by NDRI and shipped overnight on wet ice in DMEM+PSG. All specimens were received in the laboratory less than 24 hours post-surgery and were kept at 4°C throughout the collection and transportation process. Lymphocytes were extracted by dissection and maceration of the tissue in RPMI media. Lymphocyte-containing media was passed through a 40*μ*m filter and pelleted at 1500rpm for 5 minutes. RBC were lysed for 5 minutes in sterile RBC lysing solution (0.15M ammonium chloride, 10mM potassium bicarbonate, 0.1M EDTA). After dilution to 50ml with PBS, lymphocytes were counted, and pelleted. Aliquots of 5(10)^7^ to 1(10)^8^ cells were resuspended in 1ml of freezing media containing 90% FBS and 10% DMSO and cryopreserved until needed for experiments.

### Infection of primary lymphocytes with KSHV

Lymphocytes were thawed rapidly at 37°C, diluted dropwise to 5ml with RPMI and pelleted. Pellets were resuspended in 1ml RPMI+20%FBS+100*μ*g/ml DNaseI+ Primocin 100*μ*g/ml and allowed to recover in a low-binding 24 well plate for 2 hours at 37°C, 5% CO_2_. After recovery, total lymphocytes were counted and Naïve B cells were isolated using Mojosort™ Naïve B cell isolation beads (Biolegend 480068) or Naïve B cell Isolation Kit II (Miltenyi 130-091-150) according to manufacturer instructions. Bound cells (non-naïve B and other lymphocytes) were retained and kept at 37°C in RPMI+20% FBS+ Primocin 100*μ*g/ml during the initial infection process. 1(10)^6^ Isolated naïve B cells were infected with iSLK-derived KSHV.219 (dose equivalent to the ID20 at 3dpi on human fibroblasts) or Mock infected in 400ul of total of virus + serum free RPMI in 12×75mm round bottom tubes via spinoculation at 1000rpm for 30 minutes at 4°C followed by incubation at 37°C for an additional 30 minutes. Following infection, cells were plated on irradiated CDW32 feeder cells in a 48 well plate, reserved bound cell fractions were added back to the infected cell cultures, and FBS and Primocin (Invivogen) were added to final concentrations of 20% and 100*μ*g/ml, respectively. Cultures were incubated at 37°C, 5% CO_2_ for the duration of the experiment. At 3 days post-infection, cells were harvested for analysis by flow cytometry.

### Flow cytometry staining and analysis of KSHV infected tonsil lymphocytes

A proportion of lymphocyte cultures at baseline or 3dpi representing ∼500,000 cells were pelleted at 1400 rpm for 3 minutes into 96-well round bottom plates. Cells were resuspended in 100*μ*l PBS containing (0.4ng/ml) fixable viability stain (BD Cat# 564406) and incubated on ice for 15 minutes. Cells were pelleted and resuspended in 100*μ*l cold PBS without calcium and magnesium containing 2% FBS,0.5% BSA (FACS Block) and incubated on ice for 10 minutes after which 100*μ*l cold PBS containing 0.5% BSA and 0.1% Sodium Azide (FACS Wash) was added. Cells were pelleted and resuspended in FACS Wash containing B cell phenotype panel as follows for 15 minutes on ice: (volumes indicated were routinely used for up to 0.5(10)^6 cells and were based on titration of the individual antibodies on primary tonsil lymphocyte specimens) CD19-PerCPCy5.5 (2.5*μ*l, BD 561295), CD20-PE-Cy7 (2.5ul, BD 560735), CD38-APC (10*μ*l, BD 555462), IgD-APC-H7 (2.5*μ*l, BD 561305), CD138-v450 (2.5*μ*l, BD 562098), CD27-PE (10*μ*l BD 555441). After incubation, 150*μ*l FACS Wash was added and pelleted lymphocytes were washed with a further 200*μ*l of FACS Wash prior to being resuspended in 200*μ*l FACS Wash for analysis. Data was acquired on a BD FACS VERSE Flow Cytometer and analyzed using FlowJo software. Readers should note that the BD FACS VERSE analysis instrument lacks a 561nm laser so RFP lytic reporter expression from the KSHV.219 genome is not detectable in the PE channel. For baseline T cell frequencies 0.5e6 cells from baseline uninfected total lymphocyte samples were stained and analyzed as above with phenotype antibody panel as follows: CD95-APC (2*μ*l, Biolegend 305611), CCR7-PE (2*μ*l, BD 566742), CD28-PE Cy7 (2*μ*l, Biolegend 302925), CD45RO-FITC (3*μ*l, Biolegend 304204), CD45RA-PerCP Cy5.5 (2*μ*l, 304121), CD4-APC H7 (2*μ*l, BD 560158), CD19-V510 (3*μ*l, BD 562953), CD8-V450 (2.5*μ*l, BD 561426)

### KSHV neutralization via soluble Syndecan-1

Infections were performed as described above except KSHV.219 virus was pre-incubated for 30 minutes on ice with serum free RPMI only or serum free RPMI containing recombinant human syndecan-1 protein (rhCD138, BioVision, 7879-10) prior to being added to Naïve B lymphocytes. rhCD138 concentrations noted in the text indicate the final concentration of recombinant syndecan-1 in the reconstituted total lymphocyte culture. Infection was analyzed at 3 days post-infection by flow cytometry for B cell lineages and KSHV infection as detailed above. For experiments involving human fibroblasts virus was added to cells in serum free media, cells were spinoculated for 30 minutes at 1000rpm, incubated at 37°C for 1 hour, then infection media was removed and replaced with complete media. At 3dpi cells were harvested via trypsinization and analyzed for infection by flow cytometry.

### T cell depletion studies

Infections were performed as described above except a sub-population of total lymphocytes were depleted of either CD4 or CD8 T cells using positive selection magnetic beads (MojoSort™ Human CD4 T Cell Isolation Kit Cat#480009, MojoSort™ Human CD8 T Cell Isolation Kit Cat#480011). The resulting depleted fractions or unmanipulated total lymphocytes were used to reconstitute naïve B lymphocytes following infection rather than bound lymphocyte fractions as described above.

### Statistical Analysis

Data plots and statistical analysis were performed in R software{RAlanguageanden:wf} using ggplot2{ggplotElegantGra:tn} ggcorrplot{ggcorrplot:2018tg} and RColorBrewer{RColorBrewerColorB:MRqqjkfS} packages. Specific methods of statistical analysis and resulting values for significance and correlation are detailed in the corresponding figure legends.

## Results

### Variability of B cell lineages in human tonsil specimens

In order to explore the B cell lineages targeted by KSHV infection in human tonsil, we procured a cohort of 40 de-identified human tonsil specimens from donors of diverse age, sex and self-reported race (Figure 1A). Analysis of the baseline frequencies of individual B cell lineages by multi-color flow cytometry (Supplemental Figure 1A) revealed that the composition of individual human tonsil specimens is highly variable (Figure 1B). This variation was independent of donor age for many lineages. However, overall B cell frequencies declined with age as did germinal center, plasmablast and transitional B cell populations while memory and naïve populations increased in frequency with donor age (Figure 1C).

**Figure 1:**
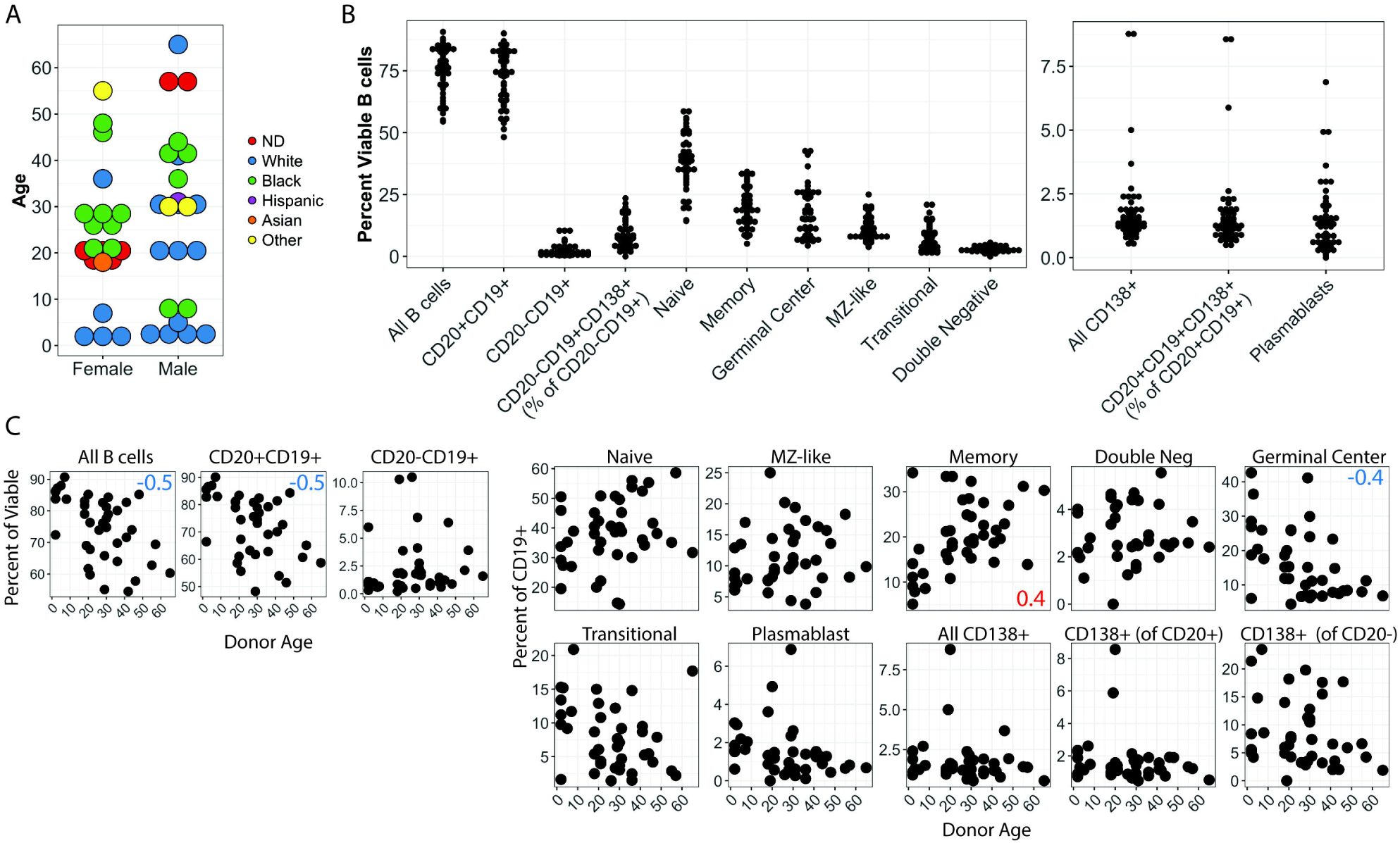
Variablility and age-dependence of B lymphocyte lineage distribution in human tonsil. (A) Donor demographics for human tonsil specimens used in this study. (B) Frequency distributions of B cell lineages in tonsil specimens (see supplemental figure 1A for lineage definitions) (C) Frequency of B cell lineages based on donor age. Pearson correlation coefficients (r) with an absolute value greater than or equal to 0.4 are shown as red or blue inset text in the subset panels.

### Variable susceptibility of tonsil-derived B cells to *ex vivo* KSHV infection

Because of the heterogeneous nature of the samples, we predicted that each sample would also have variable levels of susceptible B cell subtypes. Therefore, we employed a method for normalizing infectious dose from donor-to-donor in order to obtain cross-sectional data that was directly comparable (Figure 2A). For each sample we used magnetic sorting to isolate untouched naïve B cells, which are a known susceptible cell type {Totonchy:2018ir}, and infected 1 million naïve B cells with equivalent doses of cell-free KSHV.219 virus. After infection, bound lymphocytes from the magnetic separation were added back to each sample to reconstitute the total lymphocyte environment. Infected cultures were incubated for three days and analyzed for both B cell lineage markers and the GFP reporter present in the KSHV.219 genome to identify infected cells. Overall, susceptibility of B cells to KSHV infection varied substantially within the cohort (Figure 2B), but was not significantly correlated with age, sex or race (Figure 2C).

**Figure 2:**
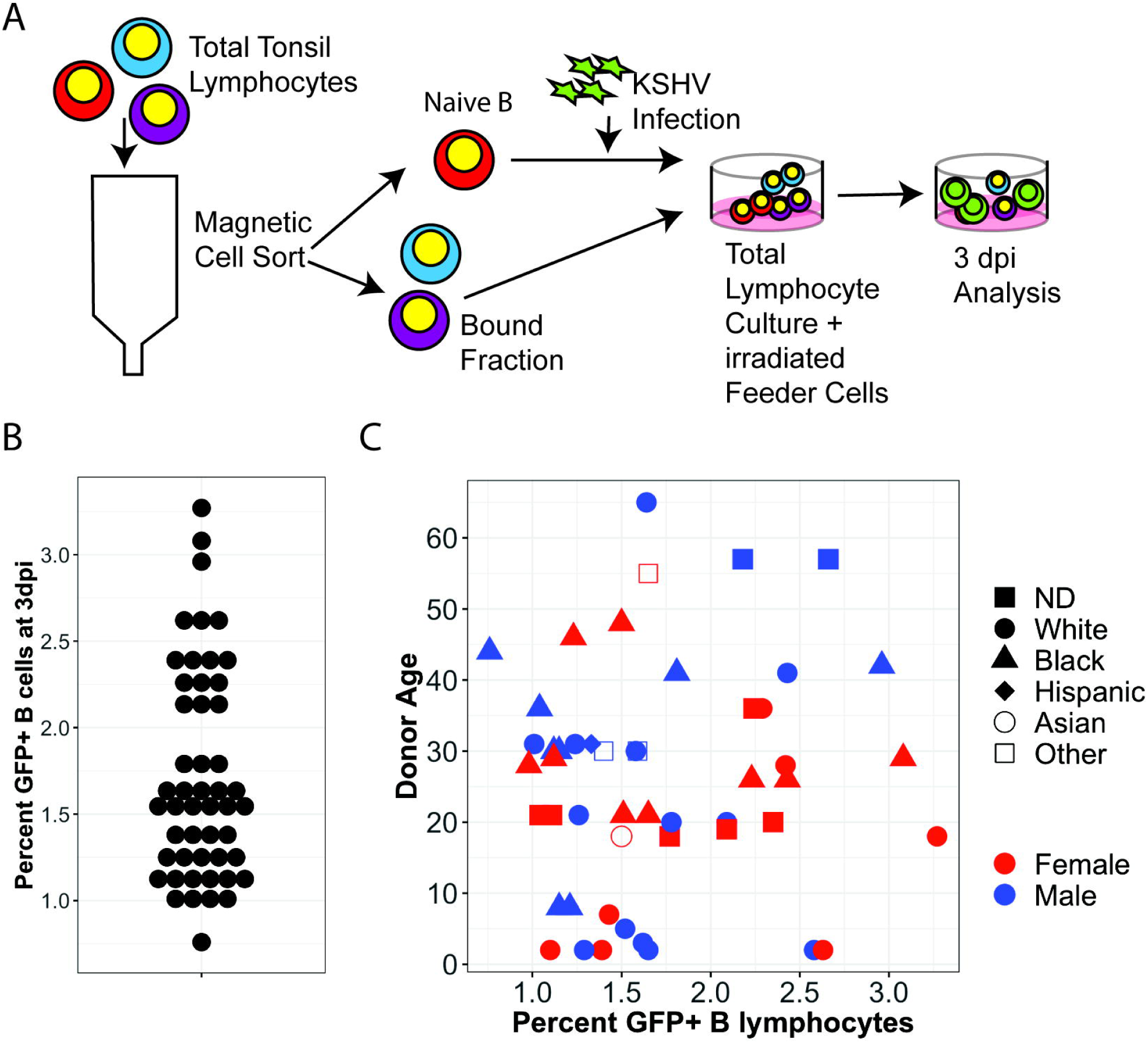
Tonsil-derived B lymphocytes from diverse donors display variable susceptibility to KSHV infection. (A) Schematic of lymphocyte infection procedure. (B) Frequency distribution for viable, CD19+GFP+ lymphocytes at 3dpi in 40 tonsil specimens. (C) Infection frequency data as in (B) displayed with respect to donor age (y-axis), sex (color), and self-reported race (shape).

### Specific targeting of individual B cell lineages by KSHV infection

We next sought to establish the B lymphocyte tropism of KSHV in the human tonsil by determining which B cell lineages are targeted for KSHV infection at early timepoints. Because levels of individual B cell lineages are highly variable between samples (Figure 1B), we represented lineage-specific susceptibility data as the percentage of GFP+ cells within each lineage so that data could be directly compared cross-sectionally within the sample cohort. Our analysis of specific B cell lineages targeted for infection by KSHV revealed that, although they represent a small proportion of the B cells within human tonsil (Figure 1B), CD138+ plasma cells are infected at high frequencies at this early timepoint. Indeed, CD19+ CD20-plasma cells displayed the highest within-lineage susceptibility of any cell type with several replicates showing 100% infection of this lineage at 3 dpi (Figure 3A). Other B cell lineages were susceptible to infection, but were infected at relatively low within-population frequencies compared to plasma cells (Figure 3A&B). Most B cell lineages showed linear correlation between within-lineage infection and overall infection, while others like plasmablast, double negative, and CD20-plasma cells showed no correlation to overall infection (Figure 3B). Pairwise correlations revealed no significant effect of the baseline (pre-infection) frequency of any B cell lineage on the susceptibility of that lineage to KSHV infection, indicating that the B lymphocyte tropism of KSHV is dependent on cell-intrinsic factors and is not simply a function of lineage population frequency (Figure 3C).

**Figure 3:**
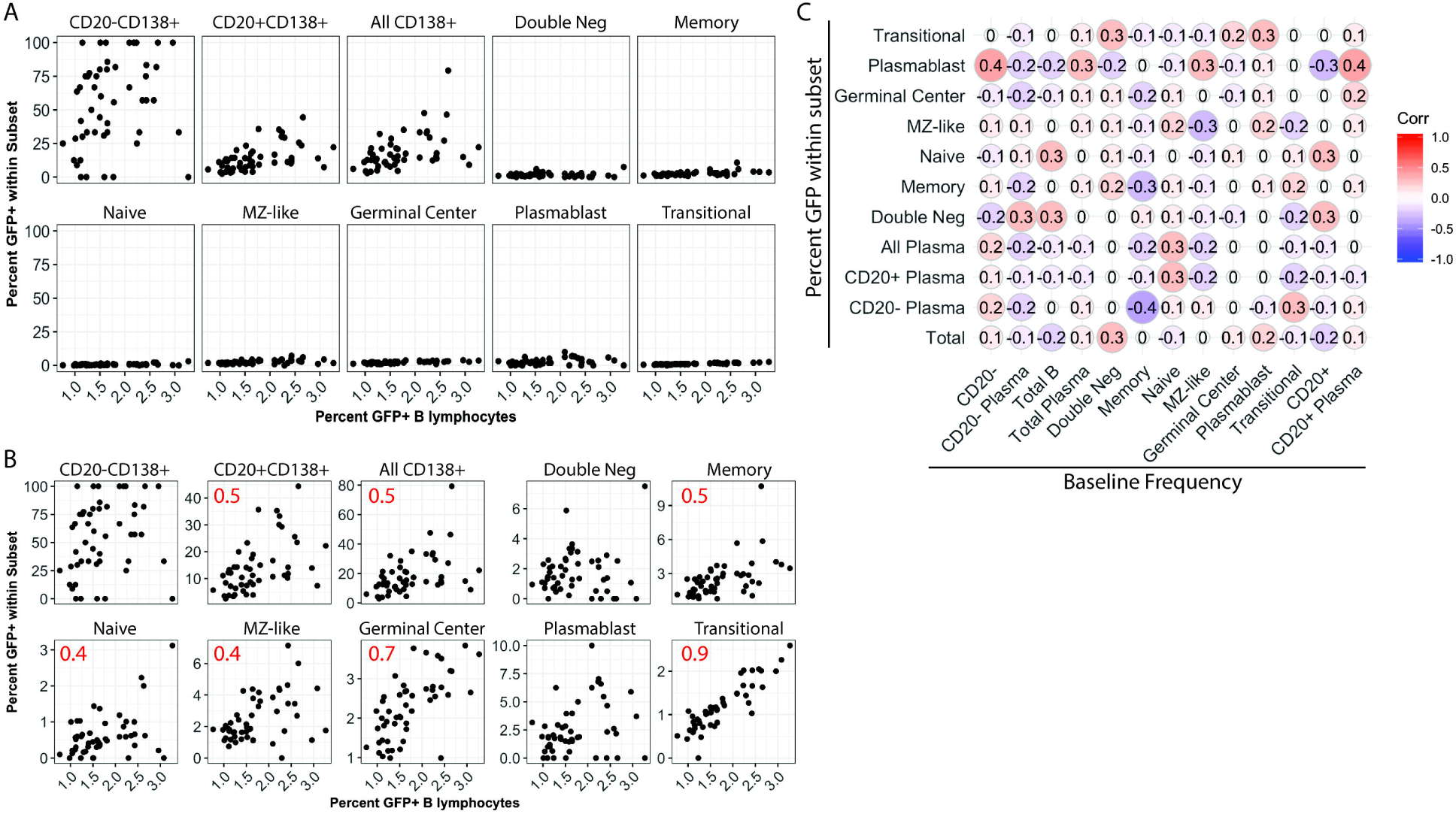
B lymphocyte lineage tropism of KSHV. Within-lineage infection frequency as a function of overall B cell infection frequency at 3dpi in for 40 tonsil specimens shown (A) normalized to 100% or (B) scaled to each individual lineage population. Pearson correlation coefficients (r) with an absolute value greater than or equal to 0.4 are shown in (B) as red inset text in the subset panels. (C) Correlation matrix showing linear relationships (Pearson’s correlation coefficient) between within-lineage infection frequency and baseline (pre-infection) overall frequency of B lymphocytes and their subsets.

Taken together, our data demonstrate that (1) tonsil lymphocyte specimens are immunologically heterogeneous, (2) susceptibility of tonsil lymphocytes to *ex vivo* KSHV infection is highly donor-dependent and (3) diverse B cell lineages are specifically targeted by KSHV early in infection. In particular, these data identify CD20-CD138+ plasma cells as a uniquely susceptible cell type for early KSHV infection in human tonsil.

### CD138 as an attachment factor for KSHV infection of plasma cells

Previous studies have shown that heparin sulfate proteoglycans (HSPG) of the syndecan family can serve as an attachment factor facilitating KSHV entry via interaction with the gH/gL glycoprotein complex ({Hahn:2009bv}. In order to test whether the high susceptibility of tonsil-derived plasma cells was due to increased attachment via CD138 (syndecan-1), we attempted to selectively neutralize KSHV entry by pre-treating cell-free virus particles with soluble recombinant CD138 (srCD138) protein prior to infection. These experiments revealed a small, reproducible decrease in overall KSHV infection of B lymphocytes with srCD138 treatment (Figure 4A). B cell lineage analysis revealed decreased infection of plasma cell lineages in some samples, but this decrease was highly variable with only 4 of 8 replicates showing a decrease. KSHV infection of other B cell lineages was largely unaffected by srCD138 treatment (Figure 4B). Moreover, KSHV infection of human fibroblasts was slightly decreased by srCD138 treatment in 4 of 6 replicates (Figure 4C). Taken together, these results suggest that, although srCD138 seems to weakly inhibit KSHV infection overall, the effect is not B cell specific. Although plasma cell infection was inhibited by srCD138 in some samples, the inconsistency between samples indicates that other factors exist which influence plasma cell targeting and the high susceptibility of plasma cells in our infection model cannot be explained by the use of plasma cell-expressed CD138 as an attachment factor for KSHV entry.

**Figure 4:**
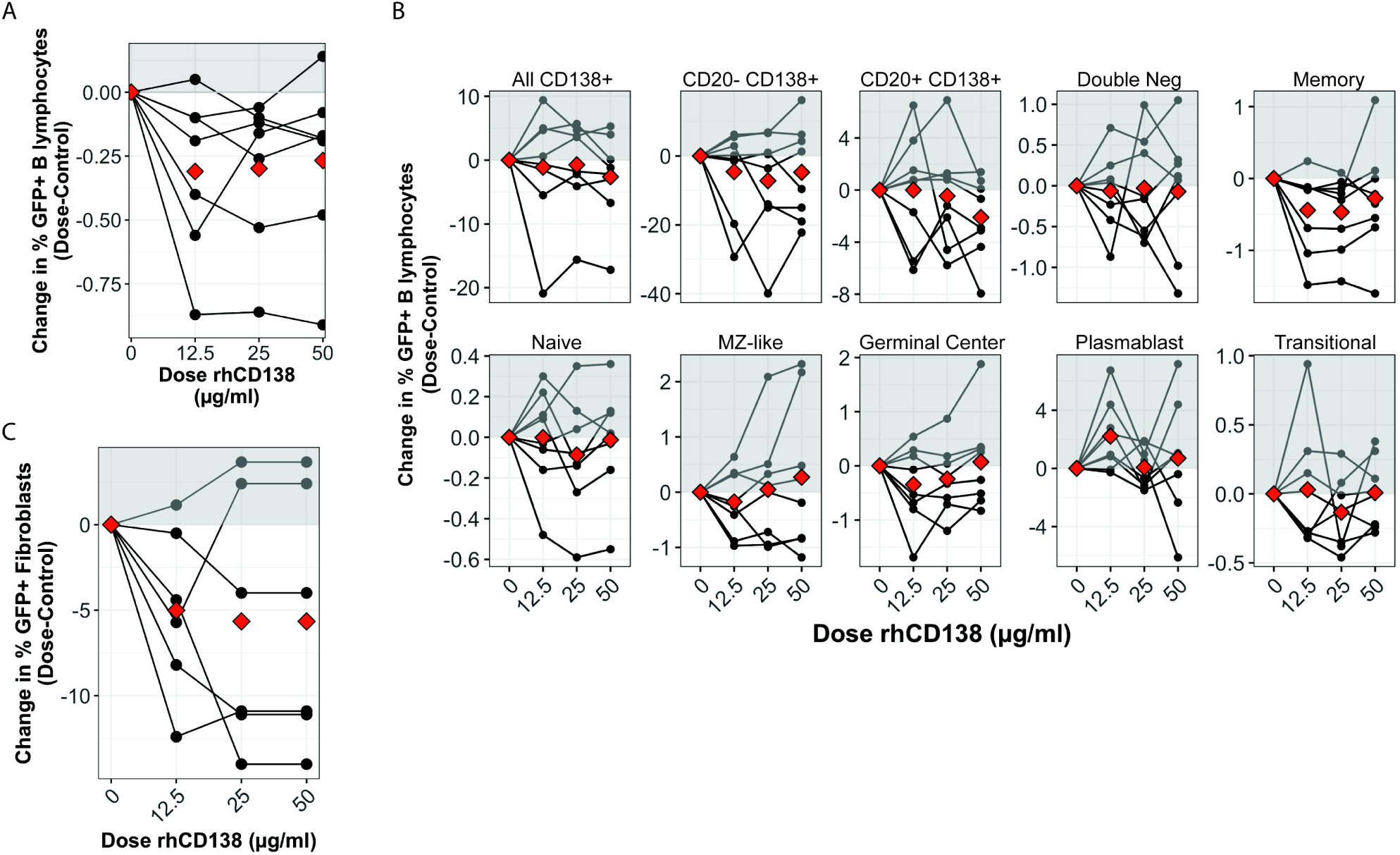
CD138 as an attachment factor for KSHV entry in plasma cells. (A) Purified KSHV virions were pre-treated with srCD138 protein and used for infection of B lymphocytes. Viable, CD19+GFP+ lymphocytes were quantitated by flow cytometry at 3dpi. 8 experimental replicates of 5 unique tonsil specimens are shown where the average infection rate was 1.8±0.5% in untreated controls. Data is represented as change in GFP+ cells at each dose of srCD138 compared to untreated control and red diamonds are the mean change at each dose. (B) Data as in (A) for within-subset GFP quantitation. (C) Data as in (A) with E6/E7 transformed fibroblasts derived from human tonsil as target cell type. 6 experimental replicates are shown where the average infection rate was 25.5±16% in untreated controls.

### Immune status alters KSHV infection of B lymphocytes

KSHV lymphoproliferative disorders occur primarily in the context of immunosuppression, and other studies have shown interactions between T cells and KSHV infected B cells in tonsil lymphocyte cultures{Myoung:2011cn}. Therefore, we wanted to determine whether the immunological composition of the tonsil lymphocyte environment would affect the establishment of KSHV infection in B lymphocytes and specifically whether overall susceptibility or targeting of particular B cell lineages is influenced by the presence or absence of T cells. Like B cell lineages, levels of CD4+ and CD8+ T cell lineages vary considerably between tonsil donors (Supplemental Figure 2A). However, unlike B lymphocyte subsets, the distribution of T cell subsets are not generally correlated with donor age (Supplemental Figure 2B). Moreover, CD4/CD8 T cell ratios are not correlated with donor age or overall susceptibility of tonsil B lymphocytes to KSHV infection (Supplemental Figure 2C).

To determine whether manipulating the T cell environment would affect KSHV infection in individual tonsil samples, we chose 11 samples with varying CD4/CD8 T cell ratios and performed KSHV infections in which total lymphocytes, CD4-depleted total lymphocytes or CD8-depleted total lymphocytes were added back following infection of sorted naïve B cells. At 3dpi, we validated T cell depletions (data not shown) and analyzed KSHV infection of B lymphocytes. Aggregated results for 11 individual tonsil specimens show that neither CD4 nor CD8 depletion significantly altered overall KSHV infection in tonsil-derived B cells (Figure 5A). However, due to the heterogeneous nature of our tonsil samples (Figure 1B & Supplemental Figure 2A), we hypothesized that the baseline T cell composition of each sample might influence the effect of T cell depletion on KSHV infection. Indeed, when the change in KSHV infection in depleted fractions is plotted against the baseline CD4/CD8 T cell ratio, we observe that KSHV infection increased when depletions were performed in samples with high baseline levels of CD4+ T cells (Figure 5B). We next analyzed the effect of T cell depletion on KSHV infection of specific B cell lineages (Figure 5C). These data reveal that depletion of CD4+ T cells increases infection of plasma cells as well as MZ-like and Transitional B cell lineages and that the effect is dependent upon the baseline CD4/CD8 T cell ratio in the sample with CD4+ T cell-rich samples showing the greatest effect. Interestingly, CD8 depletions altered infection of different lineages compared to CD4 depletions, but showed a similar dependence on the baseline level of CD4+ T cells. Taken together, these results suggest that T cells influence the dynamics of KSHV infection in the B lymphocyte compartment.

**Figure 5:**
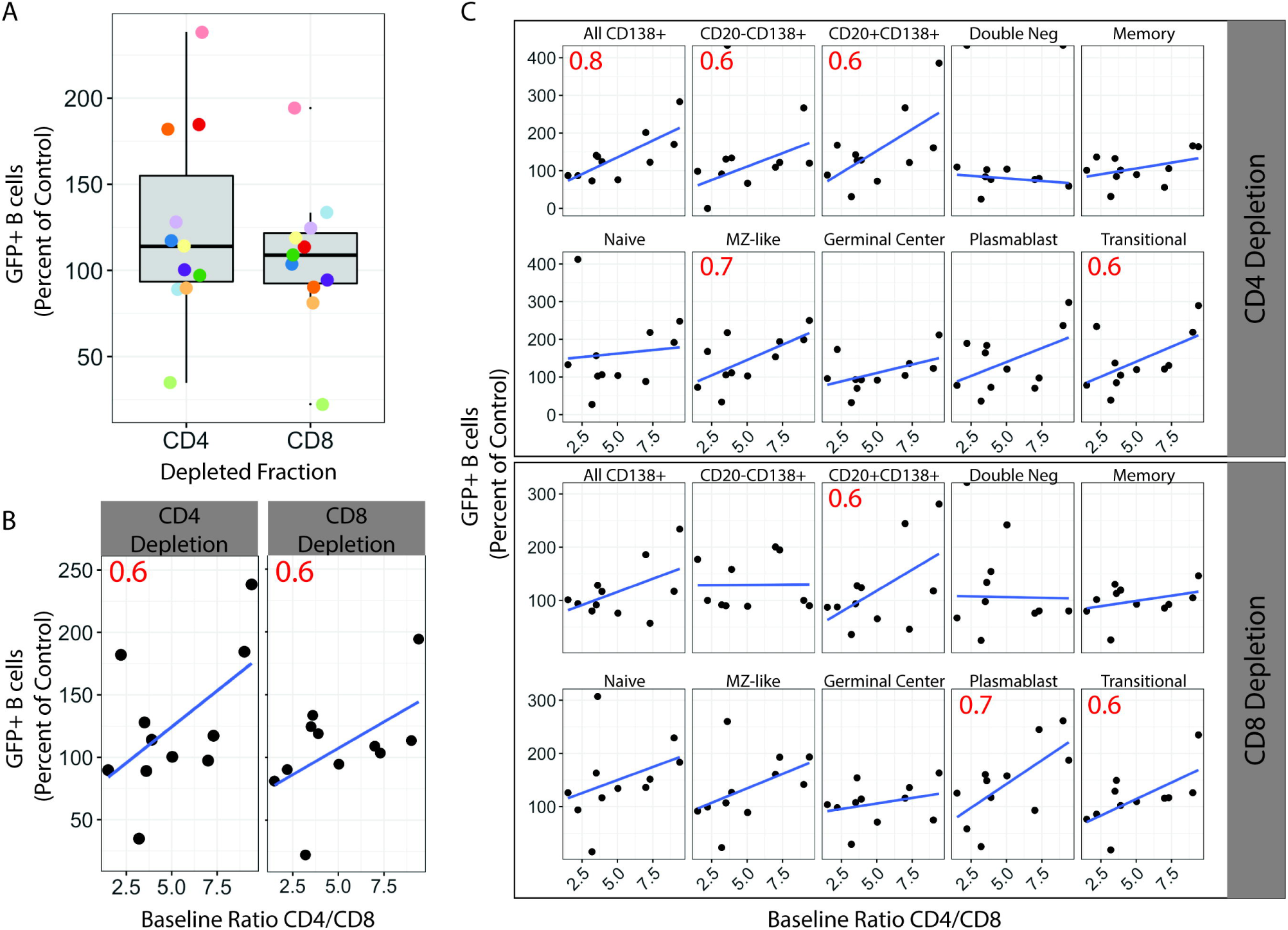
Manipulation of T cell microenvironment alters KSHV tropism for B cell lineages. Naïve B lymphocytes from 11 human tonsil specimens were infected with KSHV.219 as in Figure 2A, total lymphocytes added back following infection were either untreated or depleted of CD4+ or CD8+ T cells. Infections were analyzed at 3 days. (A) Change in GFP+ B lymphocytes in CD4 or CD8 depleted samples compared to controls. Colors indicate individual tonsil specimens. (B) infection data as in (A) plotted against the baseline CD4/CD8 T cell ratio in the sample. (C) lineage-specific infection data plotted against baseline CD4/CD8 T cell ratio. For B and C linear regression of the data is shown as a blue line and Pearson correlation coefficients (r) are shown as red text within panels only for r with an absolute value ≥ 0.6.

## Discussion

*Ex vivo* infection of tonsil lymphocytes is emerging as a viable strategy for studying early infection events for KSHV infection in B lymphocytes. However, the existing literature on KSHV infection of tonsil lymphocytes is highly varied in both approach and outcome. Hassman et. al. used cell free wild-type BCBL-1 derived KSHV virions to infect total CD19+ B lymphocytes from tonsil specimens and used staining for LANA as the only marker for infection, thus limiting their analysis to latently infected cells{Hassman:2011dj}. Bekerman et. al. also used isolated CD19+ B cells as infection targets, but employed a co-culture infection procedure using iSLK cells infected with the recombinant KSHV.219 strain employing the GFP reporter as a marker for infection. Nicol et. al. also employed a co-culture method to infect tonsil lymphocytes with KSHV.219 but used Vero cells as producers and did not isolate CD19+ B cells prior to infection. In two studies, Myoung et. al. used cell free KSHV.219 produced from Vero cells to infect mixed lymphocyte cultures. With the exception of Bekerman et. al, all of the above-mentioned studies employed some kind of activating agent (PHA stimulation or CD40L stimulation) to manipulate the activation and/or proliferation of cells *in vitro*. For our studies, we used cell-free, iSLK-derived KSHV.219 to infect Naïve B lymphocytes followed by reconstitution of the total lymphocyte environment. We also avoided activation of lymphocytes in both the isolation and culture procedures using our previously-characterized CDw32 feeder cell system {Totonchy:2018ir}. To date, no consensus has yet emerged on how to perform tonsil lymphocyte infection studies with KSHV, and how differences in infection and culture procedure influences the resulting data remains to be established.

Although previous studies in mixed lymphocyte cultures have explored limited surface markers including immunoglobulins and activation markers{Nicol:2016ga, Hassman:2011dj} and NK cell ligands{Bekerman:2013hy}, these studies essentially treated all B cells as one population. Our current study is the first to use a comprehensive panel of lineage-defining cell surface markers to carefully explore the lineage-specificity of KSHV infection in B lymphocytes.

Our finding that KSHV efficiently targets CD138+ plasma cells early in infection of tonsil B lymphocytes is particularly intriguing and relevant in the context of KSHV-mediated lymphoproliferative diseases, which often have a plasma cell or plasmablast-like phenotype{Chadburn:2008gz, Carbone:2010im}. Particularly for PEL, which uniformly presents as a clonal CD138+ neoplasm, these results suggest that the pathological cells may not be derived from KSHV-driven differentiation from less mature lineages, but instead could be the result of modifications of terminally differentiated plasma cells by direct infection. Recent studies have revealed intriguing interactions between KSHV virology and cellular mediators of the unfolded protein response (UPR) {Hu:2016gv}. The fact that the UPR is uniformly active in immunoglobulin-producing cells, like plasma cells, suggests that these cells may provide unique advantages for KSHV persistence. Certainly, our results highlight the virology of KSHV in primary plasma cells as an area urgently requiring further study.

In this study, we were unable to establish that targeting of plasma cells was due to enhanced virion binding via gH/gL interaction with the CD138 HSPG molecule, as was suggested by a previous study{Hahn:2009bv}. However, we acknowledge that selectively blocking virion binding to a specific HSPG in a primary cell system is technically difficult, and we have no way to directly verify that our neutralization of gH/gL binding sites using soluble CD138 was effective. Thus, the mechanisms underlying KSHV targeting of plasma cells and other B lymphocyte lineages for infection remains to be established. Studies in lymphoma cell lines have identified ephrin receptors as critical factors in KSHV entry{Grosskopf:2019ha, Muniraju:2019ce}. Thus, our future studies in this area will concentrate on the role of other heparin sulfate proteoglycans and Ephrin family receptors in KSHV infection of tonsillar B lymphocyte lineages.

Our characterization of immunological diversity of a large cohort of human tonsil specimens will be of interest to the general immunology community. Although a few studies have examined T cell{Petra:2015bw} or B cell{Varon:2017eo} subsets in tonsil associated with particular disease states. To our knowledge, only one other study has used multicolor flow cytometry to examine the immunological composition of both B cells and T cells in a large cohort of human tonsil samples{Stanisce:2018bc}, and this study was focused on comparing the microenvironments present in matched tonsils and adenoids rather than comparison between donors based upon demographic data as we have done here.

Our exploration of the impact of the T cell microenvironment on KSHV infection of B lymphocytes presented in this study complements previous results from Myoung et. al. showing that CD4+ T cells control lytic reactivation of KSHV in primary tonsillar B cells{Myoung:2011cn}. We make the observation that manipulating the T cell composition has a more profound effect on KSHV infection in tonsil specimens that were CD4+ T cell rich at baseline. Moreover, although the specific B cell lineages affected by depletion was different depending on whether CD4+ or CD8+ T cells were experimentally depleted, the greater effect in T cell rich samples was consistent. Thus, our data reveal that KSHV infection of B cells is sensitive to the overall immunological microenvironment and highlight that there is complexity to this relationship that cannot be adequately understood in the context of the current study. Future studies are warranted to more carefully explore the contribution of donor-specific and context-specific immunology to KSHV infection in tonsillar lymphocytes.

## Figure Legends

**Supplemental Figure 1:**
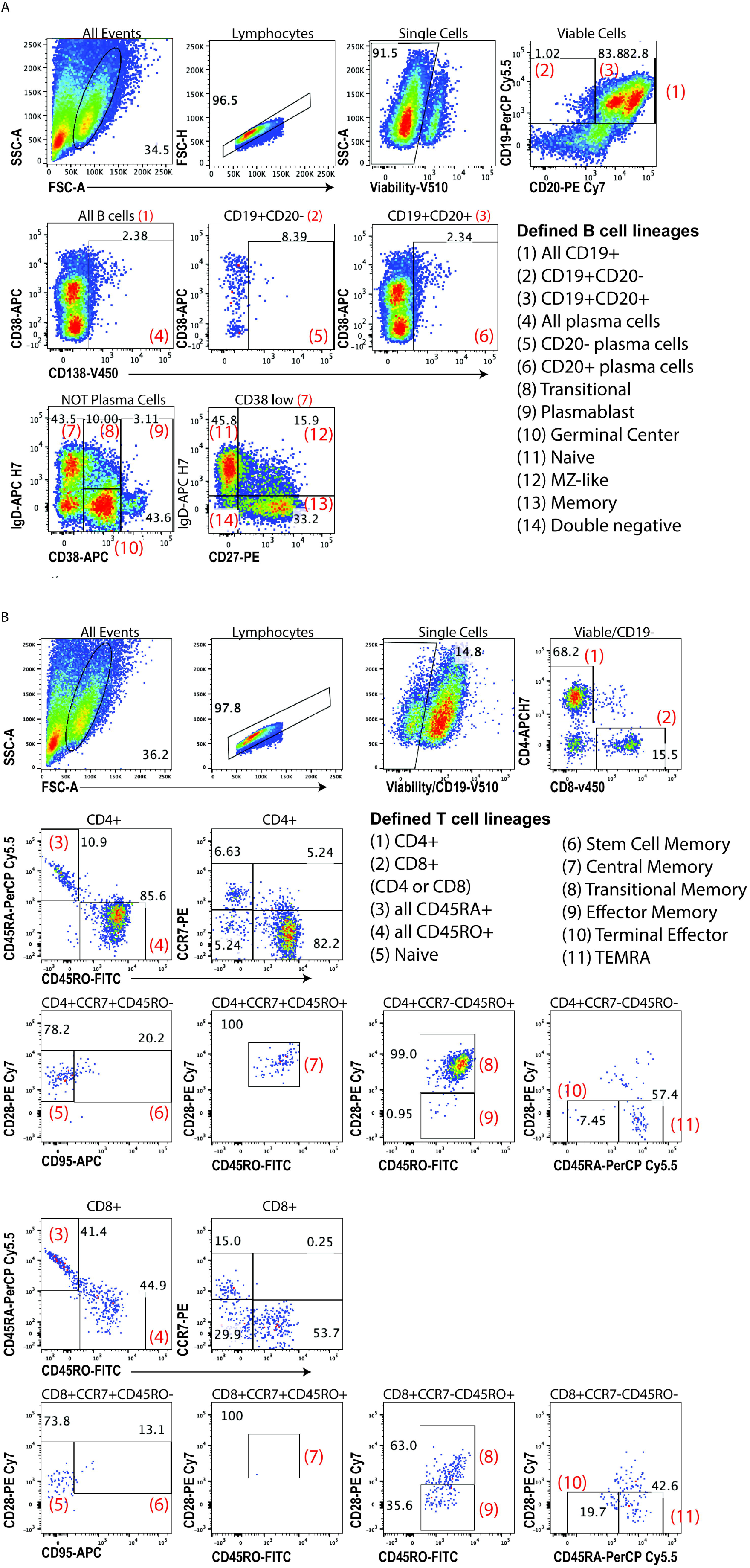
Gating schemes tonsil lymphocyte lineages. Flow cytometry data for a baseline uninfected sample from a 2-year-old male donor showing representative gating and lineage definitions used in the study for (A) B cell lineages based on {vanZelm:2007jf} and (B) T cell lineages based on {Mahnke:2013gw}.

**Supplemental Figure 2:**
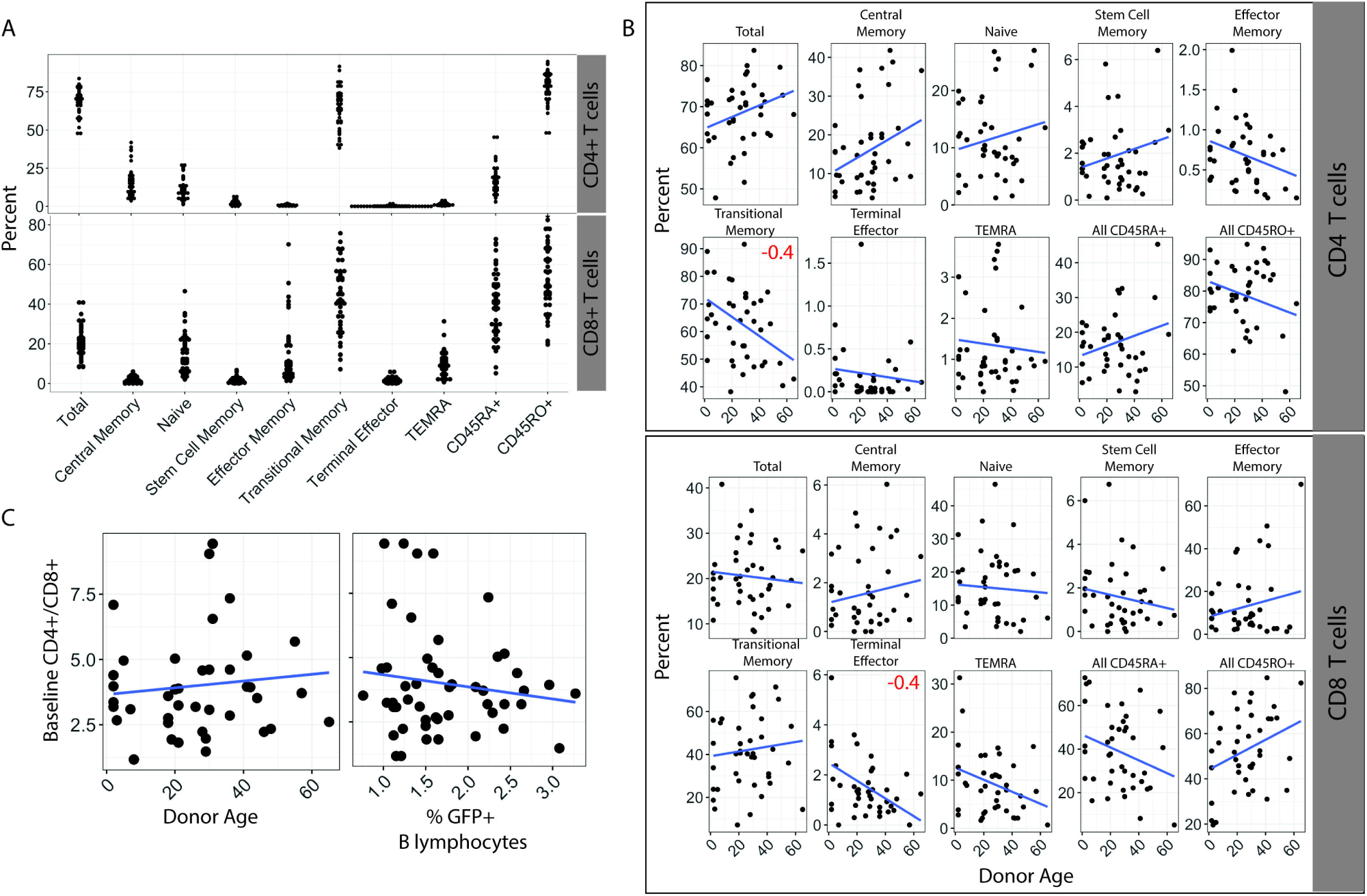
Characterization of baseline T cell microenvironment in human tonsils. (A) Frequency distributions of T cell lineages in tonsil specimens (see Supplemental Figure 1B for lineage definitions) (B) Frequency of T cell lineages based on donor age. Pearson correlation coefficients (r) with an absolute value greater than or equal to 0.4 are shown as red or blue inset text in the subset panels. (C) Correlations of Donor age (left) or B lymphocyte susceptibility (right) plotted against sample baseline CD4+/CD8+ T cell ratios.

